# A simple approximation to bias in gene-environment interaction estimates when a case might not be the case

**DOI:** 10.1101/444588

**Authors:** Iryna Lobach, Inyoung Kim, Alexander Alekseyenko, Siarhei Lobach, Li Zhang

**Affiliations:** Department of Epidemiology and Biostatistics, University of California, San Francisco, San Francisco, USA; Department of Statistics, Virginia Tech University, Blacksburg, VA, USA; Department of Public Health Sciences, Medical University of South Carolina, Charleston, SC, USA; Applied Mathematics and Computer Science Department, Belarusian State University, Minsk, Belarus; Department of Medicine, University of California, San Francisco, San Francisco, USA

**Author notes:** Corresponding author: Iryna Lobach, Ph.D., Division of Biostatistics, Department of Epidemiology and Biostatistics, University of California, San Francisco.

## Abstract

Case-control genetic association studies are often used to examine the role of the genetic basis in complex diseases, such as cancer and neurodegenerative diseases. The role of the genetic basis might vary by non-genetic (environmental) measures, what is traditionally defined as gene-environment interactions (GxE). A commonly overlooked complication is that the set of clinically diagnosed cases might be contaminated by a subset with a *nuisance* pathologic state that presents with the same symptoms as the pathologic state of interest. The genetic basis of the pathologic state of interest might differ from that of the nuisance pathologic state. Often frequencies of the pathologically defined states within the clinically diagnosed set of cases vary by the environment. We derive a simple and general approximation to bias in GxE parameter estimates when presence of the nuisance pathologic state is ignored. We then perform extensive simulation studies to show that ignoring presence of the nuisance pathologic state can result in substantial bias in GxE estimates and that the approximation we derived is reasonably accurate in finite samples. We demonstrate the applicability of the proposed approximation in a study of Alzheimer’s disease.

## INTRODUCTION

Genetic association studies that estimate the relationship between gene-environment interactions (GxE) and a complex disease have the potential to provide valuable clues to the underlying aeteology of complex diseases, such as cancer and neurodegenerative disease. Case-control studies sample a set of cases and a set of healthy controls conditionally on the disease status that is often defined based on the observed clinical diagnosis. A commonly overlooked complication is that multiple pathologic mechanisms might share symptoms and hence result in the same clinical diagnosis, i.e. the set of cases might be contaminated by a subset with a nuisance pathologic diagnosis. Frequencies of the pathologic diagnosis of interest within the set of clinically diagnosed cases might vary by the environment.

Our motivating study is a GWAS of late-onset Alzheimer’s disease (AD), a neurodegenerative disorder that is clinically characterized by progressive mental decline, but histopathologically defined by highly abundant amyloid deposits and neurofibrially tangles in the brain (Potter and Wisniewski, 2012). Recent biomarker studies of AD (Salloway and Sperling, 2015; Salloway et al, 2014) reported that 36% of ApoE *ɛ*4 non-carriers and 6% in ApoE *ɛ*4 carriers clinically diagnosed with AD do not have evidence of amyloid deposition and hence do not qualify for the *pathologic (true)* diagnosis of AD.

We are interested in estimating the association between the genetic variants serving adaptive immune system and the true, pathologically confirmed, AD status. The effect of the genetic variants might vary by the ApoE *ɛ*4 status and in the context of this study we define ApoE *ɛ*4 to be the environment. It is possible that the symptoms resulting in the clinical diagnosis are manifestations of an underlying polygenic mechanism and hence the clinical diagnosis is a surrogate of the *true* diagnosis. It is also possible that the symptoms with and without the amyloid deposition evidence represent diverse mechanisms each with a distinct genetic basis. In the latter case the usual logistic model with the clinical diagnosis as an outcome is based on a misclassified disease status and hence misspecifies the link between GxE and the AD status.

Traditionally, case-control GWAS are analyzed in a logistic regression model as if the data are collected prospectively based on a justification provided by Prentice and Pyke (1979). In the situation when the disease status is misclassified with frequencies varying by the environment, the result of Prentice and Pyke (1979) does not naturally extend.

Extensive literature (Carroll et al, 2006) reports how the estimates of the main genetic effects can be biased in situations when the disease status is misclassified. We have recently examined bias in GxE when misclassification probabilities vary by the environment and proposed a solution that alleviates the bias (Lobach et al, 2018). This solution requires optimization of a complex non-linear function. Interestingly, Neuhaus (1999) derived a general approximation to the bias in a univariate setting when the data are collected prospectively and are analyzed in the logistic regression model. We extend the literature by deriving a general theoretical bias and a convenient approximation to the bias for GxE when the data are collected retrospectively and hence in a logistic regression model where both the design of data collection and presence of the nuisance pathologic diagnosis in the set of cases are ignored.

## MATERIALS AND METHODS

We define *G* be the genotype, e.g. SNPs measured at multiple locations. Let *X* be the environmental variable that interacts with *G*. We assume that the genotype is independent of all environmental variables and the genotypes follows Hardy-Weinberg Equilibrium: *G*~*Q* (*g*, *θ*).

We define *D^CL^* = {0, 1} be observed clinical disease status defined based on a set of symptoms. Suppose that the same set of symptoms can be caused by two distinct pathophysiologic mechanisms. Let *D* be the *true* disease status defined based on the underlying pathology, where *D* = 1 indicates the disease of interest, while *D* = 1* is the nuisance disease. Thus pr(*D* = 1) + *pr*(*D* = 0) + *pr* (*D* = 1*) = 1. For ethical and/or budgetary reasons it might not be possible to measure the underlying pathology on all patients, hence *D* is latent. Instead, an evaluation might be performed on a subset of patients or in an external reliability study. We define τ(*X*) = pr(*D* = 1 |*D* ^*CL*^ = 1, *X*) to be the frequency of the *true* diagnosis of interest within the clinically diagnosed set that varies by the environmental variable %. In our setting the clinical diagnosis of healthy controls corresponds to *D* = 0, hence pr(*D* = 0| *D^CL^* = 0, *X*) = 1, pr(*D* = 0 |*D^CL^* = 1, *X*) = 0. We let probabilities of the clinical and *true* diagnoses in the population to be 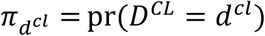 and *π_d_* = pr(*D* = *d*), respectively. Let 𝒮(*X*) = pr(*D* = 0| *D^CL^* = 0, *X*). For clarity of presentation we suppose that all variables are binary. The setting can be easily extended to accommodate multi-level categorical variables.

We first consider a binary setting where the risk parameters are defined in terms of *D* = 1 vs. *D* = 1* and *D* = 0 combined. Then the true risk model in terms of coefficients Β = (*β*_0,_ *β*_G,_ *β*_X,_ *β*_G×X_) is

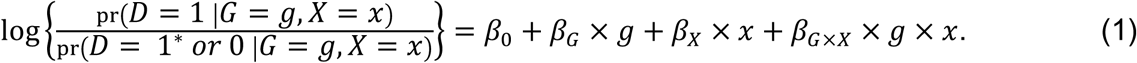

In the second setting we consider the risk model is defined separately for *D*= 1 vs. *D* = 0 in terms of Β = (*β*_0,_ *β*_G,_ *β*_X,_ *β*_G×X_) and for *D* = 1* vs. *D* = 0 in terms of 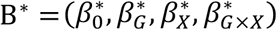 by

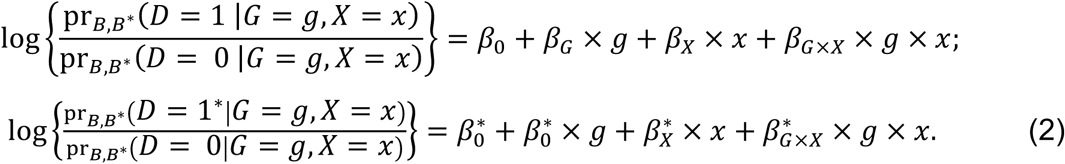

In equation (2) Β and Β* might share coefficients, e.g. if 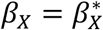. While we define the parameters of interest to be Β = (*β*_0,_ *β*_G,_ *β*_X,_ *β*_G×X_), these parameters are independent and are different in model (1) and model (2).

Suppose that the clinical-pathological diagnoses relationship is ignored and the clinical diagnosis is used as the outcome variable. Then the risk model is

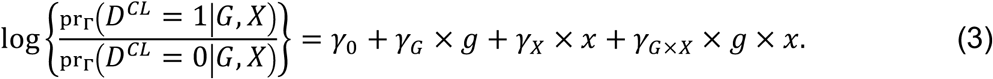

Derivations provided in Online Methods show that

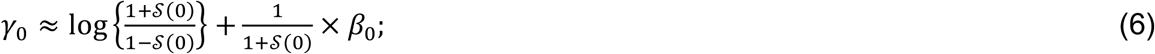

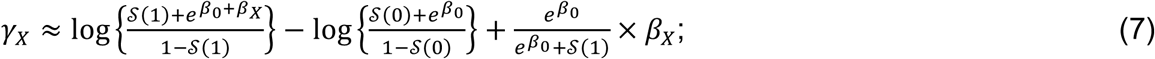

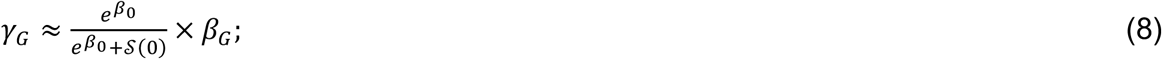

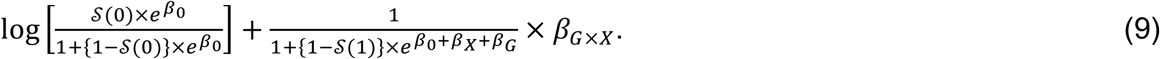

<u>Remarks on formulas (6)-(9):</u>

1. Online Methods section provides formulas (A11)-(A15) for the setting with environmental variable Z that does not interact with the genetics.
2. When the clinical diagnosis and pathologic disease status correspond, i.e. 𝒮(*X*) = 0 for *X* = 1 and *X* = 0, then all parameter estimates are unbiased
3. If *β_G_* = 0, then *γ_G_* = 0. Hence the usual logistic regression yields consistent estimate of the null *β_G_*.
4. *β*_0_ = 0 does not necessary results in *γ*_0_ = 0. Similarly, if *β_X_* = 0, then *γ*_X_ might not be zero and if *β*_G×X_ = 0, then γ_*G* ×*X*_ might not be zero. Hence the usual logistic regression does not yield consistent estimate of the null effect *β*_0,_ *β*_G,_ *β*_X,_ *β*_G×X_.
5. If *β*_L_ = 0 and *β*_G×X_ = 0 then γ_*G*_ = 0 and γ_G×X_ = 0. Hence the usual logistic regression yields consistent estimate of the null *β*_G_ and *β_G_*_×*X*_.
6. If the misclassification is non-differential, i.e. 𝒮(0) = 𝒮(1); then if *β*_X_ = 0, then γ_*X*_ = 0. That is then the usual logistic regression model yields consistent estimate of the null effect *β_X._*
7. If the misclassification is non-differential, i.e. 𝒮(0) = 𝒮(1); then if *β*_0_ = 0, *β_X_* = 0, *β_G_* = 0, *β_X_*_×*G*_ = 0 then γ_*G×X*_ = 0. That is then the usual logistic regression model yields consistent estimate of the null effect of *β_G_*_×*X*_.

We next suppose that the true model is (2), while the parameters are estimated based on a misspecified model (3).

Derivations provided in Appendix show that

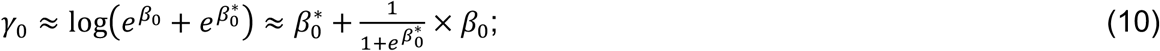

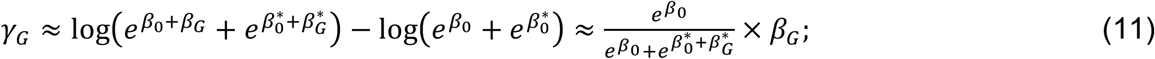

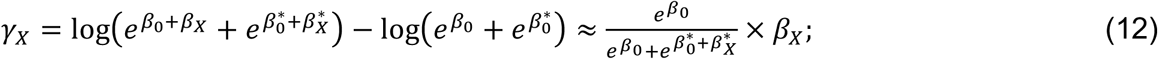

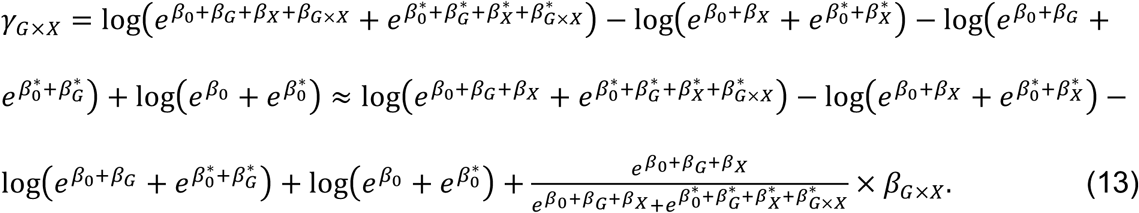

<u>Remarks on formulas (10)-(13):</u>

1. Online Methods provides formulas (A19)-(A23) for the setting with environmental variable t that does not interact with the genetics.
2. If *β*_0_ = 0 and 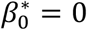, then γ_0_ = 0. That is then the usual logistic regression yields the consistent estimate of the null *β*_0._
3. If *β*_G_ = 0 and 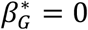 then *γ_G_* = 0. Hence in this case the usual logistic regression yields the consistent estimate of the null *β*_G_.
4. If *β*_X_ = 0 and 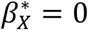, then γ_X_ = 0. Hence in this case the usual logistic regression yields the consistent estimate of the null *β_x_.*
5. If *β*_G_ = 0 and *β_X×_*_G_ = 0 then 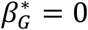 and 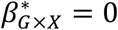. Hence the usual logistic regression yields the consistent estimate of the null *β*_G_ and *β_X×_*_G_.
6. If *β*_G_ = 0, 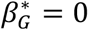, *β*_X_ = 0, 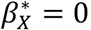 *β*_G×X_ = 0 and 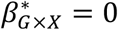, then γ_G×X_ = 0. Hence if all these parameters are zero, the usual logistic regression yields the consistent estimate of the null *β_X×G_*.

Online Methods section describes extensive simulation studies where we examined performance of the approximation empirically.

### Simulation Experiments

We perform a series of simulation experiments to evaluate the accuracy of the proposed approximation. We set our other parameters to be similar to the values observed in our GWAS of AD. We let the genotype (*G*) be a Bernoulli random variable with frequency of 0.10 to mimic a SNP and allow its effect to follow a recessive or dominant model. We simulated age (*Z*_1_) to be Bernoulli with frequency of 0.50 e.g. corresponding to a median split. Sex (*Z*_2_) is Bernoulli with *pr*(*Z*_2_=1)=0.52. The binary variable *X* = {*ɛ*4+, *ɛ*4−}, which represents the ApoE *ɛ*4 status according to presence or absence of !4 allele that occurs in approximately 14% of the population. Similarly to Salloway, et al (2014), we define the proportion of the nuisance disease within the clinical diagnosis is defined as *pr*(*D*=1^*^|*D^CL^*=1, *ɛ*4–)=0.36 and *pr*(*D*=1^*^|*D^cl^*=1, *ɛ*4+)=0.06.

#### Setting A

We first simulate data according to model (1) while estimate parameters in model (3) where the clinically diagnosed status is the outcome variable. The disease status is simulated according to model (1) with β_*G*×*ɛ*4_ varying from log(1) to log(8) and we set 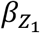, to be 0, 0.5, 1, 1.5. The other risk coefficients are *β*_0_ = −1, *β_G_* = 0.41, 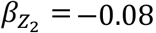, 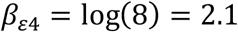. We simulated 500 datasets with 3,000 cases and 3,000 controls. Shown in **Supplementary Table 1** are parameter estimates when *β*_*G*×*ɛ*4_ = 0 obtained as an average (empirical estimate) and standard deviation (SD) across the 500 simulated datasets, theoretical values and the simple approximation derived in (A11)-(A15). For all parameters the empirical values are close to the approximation with the average difference across all parameters is 0.11. The approximation was furthest away from the empirical estimate of main effect of ApoE *ɛ*4 status, with the empirical estimate being 1.80, while the approximation is 1.30.

Shown on **Supplementary Figures 1A-D** are the estimates (empirical estimate that is the average across 500 simulated datasets (AVE), theoretical estimate (TH) (A11)- (A15) and approximation (APX) (A11.1)-(A15.1) across the levels of 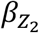 that are indicated by color and across values of *β*_*G*×*ɛ*4_ across panels of each graph. In all of these settings the theoretical values and the approximations are accurate relative to the empirical estimates. The furthest from the empirical estimates are approximation for the main effect of ApoE *ɛ*4 status with the difference mainly driven by the approximation.

To examine robustness of the theoretically derived magnitude of bias to misspecification of proportions of the nuisance disease, we calculated the theoretical values while underestimating the frequencies to be *pr* (*D* =1^*^|*D^C^*^L^ =1, *ɛ*4 –) =0.30 and *pr* (*D* =1^*^|*D^cl^* =1, *ɛ*4 +) =0 (**Supplementary Figure 2A-C** for *γ_G_*,*γ ɛ*_4_, *γ*_*G*×_*ɛ*_4_). The approximation to bias is robust to this misspecification and while overestimating the frequencies to be *pr* (*D* =1^*^|*D^C^*^L^ =1, *ɛ*4 –) =0.42 and *pr* (*D* =1^*^|*D^cl^* =1, *ɛ*4 +) =0.12 (**Supplementary Figure 3A-C** for *γ_G_*,*γ ɛ*_4_, *γ*_*G*×_*ɛ*_4_).

#### Setting B

We next simulate the data according to the risk model (2) while estimating parameters based on the model (3). We simulate the disease status *D*=1 vs. *D*=0 based on parameters *β*_0_= −1, *β_G_*= −0.69, 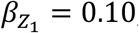, 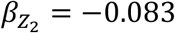, *β*_*ɛ*4_ = 1.3, *β*_*G*×*ɛ*4_ = 1.099; and we simulate *D*=1* vs. *D*=0 using 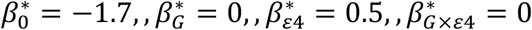 with main effects of *Z*_1_ and *Z*_2_ that are the same as for *D*=1 vs. *D*=0. With these parameters, the frequencies of the disease of interest and the nuisance disease are pr(*D* =1)=25.1%, pr(*D* = 1*) = 12.5%, pr(*D* = 1| *ɛ*4 +) = 45.4%, pr(D = 1* |*ɛ*4 +) = 16.1%, pr(D = 1|*ɛ*4 −) = 20%, pr(D = 1*|*ɛ*4 −) = 16.1%. **Supplementary Table 2** (*n*_0_ = *n*_1_= 3,000) presents empirical estimates, theoretical values (A16)-(A22) and approximation (A16.1)-(A22.1).

We first note that when presence of the nuisance disease is ignored, estimates of *β*_0_, *β*_*ɛ*4_, *β*_*G*×*ɛ*4_, *β*_G_ are substantially biased. The approximation that we derived is accurate relative to the empirical averages of the parameter estimates. For example, empirical estimate of γ_*ɛ*4_ is 1.08, while the approximation is 1.08. Empirical estimate of *γ*_*G*×*ɛ*4_ is - 0.10, while the approximation is −0.05.

Shown on **Supplementary Figures 4A-B** are the estimates (empirical estimate that is the average across 500 simulated datasets (AVE), approximation (A20)-(A23) across values of 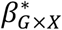 along the x-axis when *β*_*G*×*ɛ*4_= 0 (A) and when *β*_*G*×*ɛ*4_= −0.9 (B). In all of these settings the theoretical values and the approximations are accurate relative to the empirical estimates.

#### Analyses of Genetic Variants Serving Adaptive Immune System in Alzheimer’s Disease

We applied the proposed analyses to a dataset collected as part of the Alzhemer’s Disease Genetics Consortium. The data consists of 1,245 controls and 2,785 cases. The average age (SD) of Cases and controls are 72.1 (9.1) and 70.9 (8.8) years, respectively. Among cases, 1,458 (52.4%) are men; among controls, 678 (63.9%) are men. At least one ApoE *ɛ*4 allele is present in (64.5%) of cases and 365 (29.1%) of controls.

Illumina Human 660K markers have been mapped onto human chromosomes using NCBI dbSNP database (https://www.ncbi.nlm.nih.gov/projects/SNP/). Chromosome location, proximal gene or genes and gene structure location (e.g. intron, exon, intergenic, UTR) has been recorded for all SNPs. We inferred the adaptive immune system pathways based on the information from Kyoto Encyclopedia of Genes and Genomes (https://www.genome.jp/kegg) (KEGG), gene ontology (GO) consortium (https://www.geneontology.org) and Ariadne Genomics (https://www.ariadnegenomics.com). From these data with quality control measures (observed frequency of minor allele >5%), we inferred 133 SNPs to reside in genes serving the adaptive immune system.

It is of interest to examine a relationship between the pathologic diagnosis and each of the 133 SNPs (*G*), ApoE *ɛ*4 status (*X*), age (*Z*_1_), sex (*Z*_2_). The effect of SNPs might vary by ApoE *ɛ*4 hence we included interaction between the genotype and ApoE *ɛ*4 status.The genetic variables are modeled using a binary indicator of presence or absence of a minor allele.

We estimate parameters using the standard logistic model (3) that uses the clinical diagnosis as a surrogate of the pathophysiologic. We assume that the proportion of nuisance disease is as estimated by Salloway (2015) study, i.e. the proportion of the nuisance disease within the clinically diagnosed set of cases is 36% in ApoE *ɛ*4 non-carriers and 6% in ApoE *ɛ*4 carriers. We first assume that the true model is (1), i.e. the susceptibility model is defined for amyloid-related AD symptoms vs. healthy controls and non-amyloid AD symptoms combined. In this setting we estimate the magnitude of bias using approximation (A11.1-A15.1). We next assume that the true model is (2), i.e. the susceptibility model is defined for amyloid-related AD symptoms vs. controls and for non-amyloid-related symptoms vs. controls. We then estimate the magnitude of bias using (A19.1)-(A25.1).

Shown on **Figures 1 and 2** are the estimated biases in the main effect of each SNP, ApoE *ɛ*4 4 and interaction between the SNPs and ApoE *ɛ*4 4 status. We first note that the bias in the estimate can be substantial. For example, in the usual logistic regression with the clinical diagnosis as an outcome variable, main effect of rs597587 is statistically significant with Bonferroni correction (p<0.05/133). The estimate is 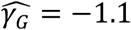, while if model (1) is the true model, the bias is approximated to be 2.9 and the interactive effect is estimated to be 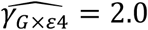, *p* = 0.008, which means if the model was incorrectly specified, both main and interactive effects would be mistakenly interpreted. For another SNP, rs12111032, main effect estimate is 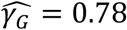 with p<0.05/133, while the bias is −0.04 if model (1) is the true model and −0.17 if model (2) is the true model.

**Figure 1:**
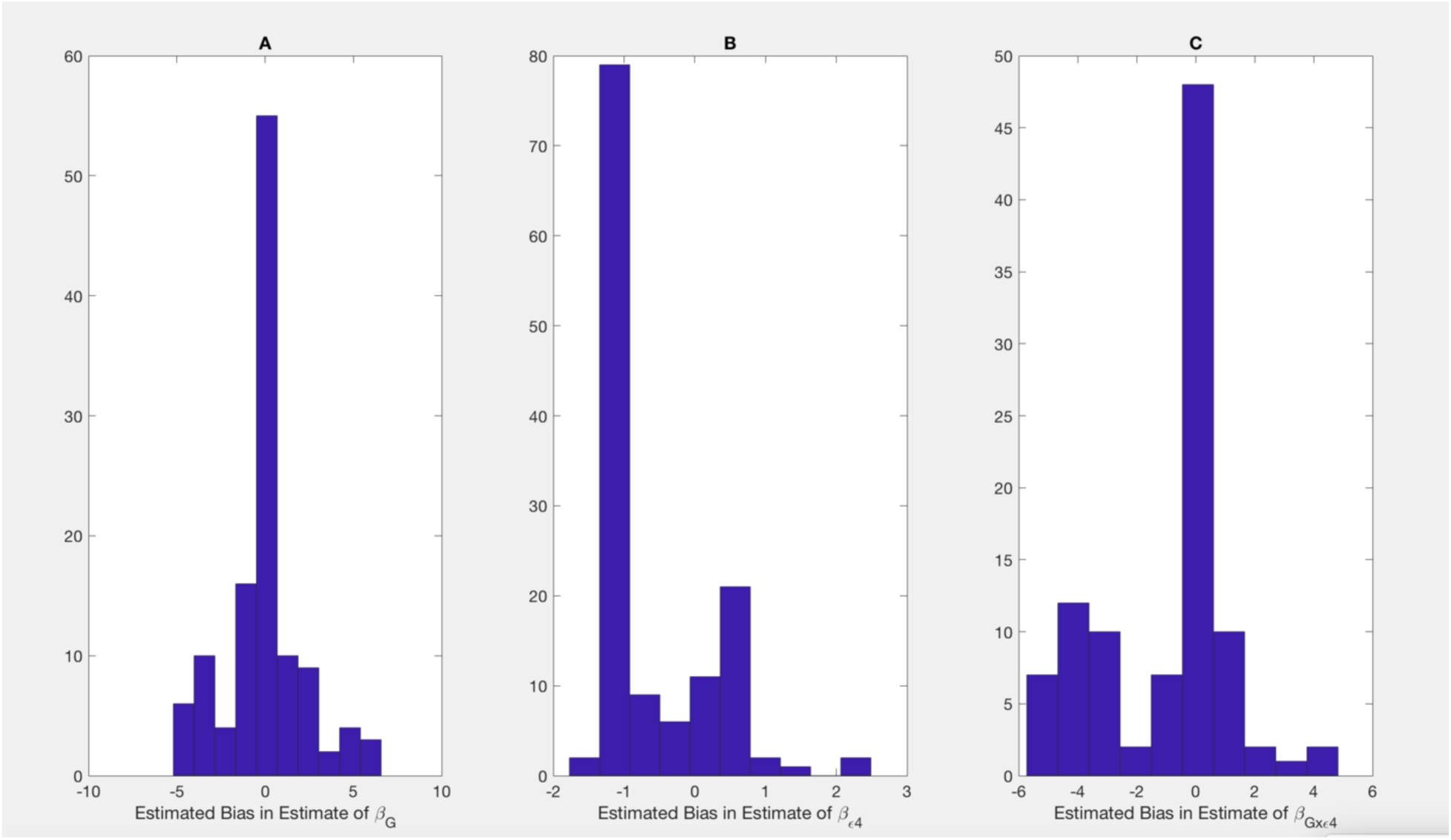
Bias in parameter estimates in Alzheimer’s disease study assuming that the true model is (1) while parameters are estimated using model (3). Magnitude of bias is approximated using (A11.1)-(A15.1).

**Figure 2:**
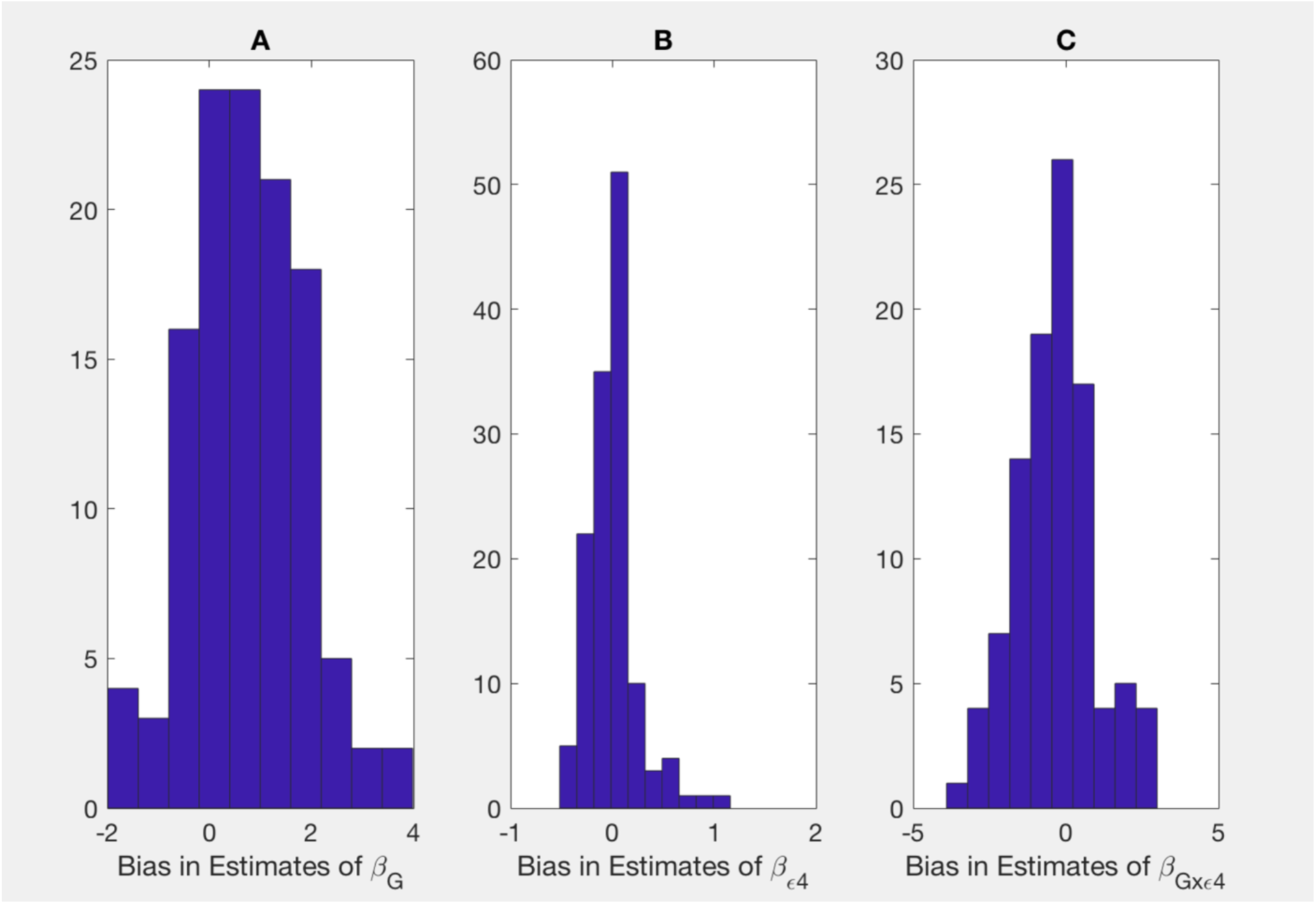
Bias in parameter estimates in Alzheimer’s disease study assuming that the true model is (2) while parameters are estimated using model (3). Magnitude of bias is approximated using (A19.1)-(A23.1).

**Table 1** presents estimates of main effects of SNPs and p-values obtained using the usual logistic regression model with the clinically diagnosed disease status as an outcome variable for SNPs with p-value <0.05 for 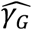 or 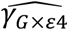 We also mapped the genes from Table 3 to the amyloid pathway using KEGG and GO. SNP, rs10059242, residing in an intergenic region between HTR4 and ADRB2 (ADRB2 is found to be in the amyloid pathway), has the main effect of 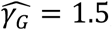, *p* = 0.04, with bias ≈ −0.9 (1), 1.8 (2) and the GxE interaction of 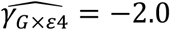, *p* = 0.04 with bias ≈ −0.9 (1), −2 (2).

**Table 1:** Estimate of main effect of SNPs 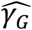 and their interaction effect with ApoE *ɛ*4 status in the Alzheimer’s disease study. The estimates and p-values are obtained using the usual logistic regression model with the clinically diagnosed status as an outcome variable (model 3). Magnitude of bias is estimated using approximation (A11.1-A15.1) assuming the true model is (1) and using approximation (A19.1)-(A25.1) assuming the true model is (2). Boldfaced are the results with p-value<0.05/13.

| SNP | Gene/Intergenic Region | Bias in $\widehat{\gamma}_G$ | | $\widehat{\gamma}_G$ from model (3) | | Bias in $\widehat{\gamma}_{G \times \varepsilon 4}$ | | $\widehat{\gamma}_{G \times \varepsilon 4}$ from model (3) | |
| --- | --- | --- | --- | --- | --- | --- | --- | --- | --- |
|  |  | (1) is<br>the<br>true<br>model | (2) is<br>the<br>true<br>model | Estimate | p-value | (1) is<br>the true<br>model | (2) is<br>the true<br>model | Estimate | p-value |
| rs401904 | CD1D \| CD1A | 0.08 | -0.05 | -0.08 | 0.64 | -0.08 | 0.25 | 0.83 | 0.04 |
| rs1748383 | N4BP2 \| RHOH | 0.07 | -0.39 | -0.07 | 0.69 | -0.07 | -0.68 | 0.82 | 0.04 |
| rs13386118 | CXCR4 \| THSD7B | 3.9 | 1.4 | -3.6 | 0.003 | 2.9 | -1.2 | 0 | 0.99 |
| rs12692222 | LOC100419686\|<br>LOC151171 | 0.04 | -0.06 | 0.01 | 0.96 | -0.05 | 0 | 1.3 | 0.04 |
| rs1645732 | FYB \| C9 | -0.21 | 0.56 | 1.1 | 0.0496 | 0.21 | -4.5 | -1.5 | 0.06 |
| rs10059242 | HTR4 \| ADRB2 | -0.90 | 1.8 | 1.5 | 0.04 | 0.89 | -2.1 | -2.0 | 0.04 |
| rs12111032 | HLA-C \| HLA-B | -0.04 | -0.17 | **0.78** | **<0.0001** | 0.04 | 0.23 | -0.27 | 0.45 |
| rs9275383 | HLA-DQB1 \| HLA-DQA2 | -0.28 | 0.78 | 2.3 | 0.002 | NA | 4 | NA | 0.99 |
| rs2551698 | GSR | 3.4 | 0.07 | -0.60 | 0.003 | -4.5 | 0.41 | 0.32 | 0.43 |
| rs12543466 | ANGPT1 \| RSPO2 | -0.17 | 0.84 | 0.53 | 0.26 | 0.18 | -1 | -1.6 | 0.03 |
| rs597587 | MYEOV \| CCND1 | 2.9 | -0.25 | **-1.1** | **<0.0001** | -2.9 | 1.0 | 2.0 | 0.008 |
| rs1586910 | DCN \| BTG1 | 0.12 | -0.75 | -0.14 | 0.38 | -0.13 | 0.64 | 1.1 | 0.006 |
| rs6018027 | SRC | 2.7 | -1.8 | -0.26 | 0.08 | -3.4 | 0.35 | 0.65 | 0.04 |
| rs4969754 | RPS6KA3 \| CNKSR2 | 6.6 | 0.29 | -0.53 | 0.02 | -4.9 | -0.09 | 0.03 | 0.94 |

For three SNPs, rs401904, rs1748383, and rs12692222, the estimates in the usual logistic model (3) are nearly zero, while biases estimated assuming model (1) is true are nearly zero as well. This is consistent with the theoretical observations that the null effect estimate based on the logistic regression can be unbiased.

## DISCUSSION

We derived a simple and general approximation to bias in GxE parameter estimates when multiple pathologic mechanisms present with the same set of symptoms. The approximation to the bias relies on the estimates of the frequencies of the pathologic disease state within the clinical diagnosis. The approximation that we derived complements a recent study (Lobach et al, 2018) where we developed a pseudolikelihood method to incorporate uncertainty about clinical-pathological diagnoses relationship, where the solution requires optimization of a complex non-linear function. The approximation that we derived provides a simple formula that is intuitive and easy to apply.

We observed that parameter estimates could be substantially biased when presence of the nuisance pathology is ignored. This observation has been also made by Manchia et al (2013), where the simulation studies and data analyses on the main effect of genotype showed that the risk parameter estimates attributable to the genetics could be largely underestimated.

In our analysis of Alzheimer’s disease the reliability study is based on 1,121 carriers of ApoE *ɛ*4 allele and 1,331 non-carriers of ApoE *ɛ*4 allele. Hence we suppose that the clinical-pathological diagnoses relationship is estimated reasonably reliably well. Moreover, the reliability study is performed on the same patient population, i.e. patients followed-up by the Alzheimer’s disease centers. There is substantial, although not exactly known, overlap between the set with genotypes and the reliability set. In general, we advocate sensitivity analyses that examine potential differences in the parameter estimates due to misspecifications of frequencies of the pathologic diagnosis within the set of clinically defined cases.

While our study is motivated by a specific example of Alzheimer’s disease, the application of the approximation that we derived is readily applicable to other diseases. For example, recent studies report that the underlying biologic mechanisms of breast cancer vary by expression of progesterone and estrogen. Frequencies of subtypes can be estimated based on SEER database (https://seer.cancer.gov/).

## Data availability

The data analyzed in this work is available at the Database of Genotypes and Phenotypes, study accession number phs000372.v1.p1(https://www.ncbi.nlm.nih.gov/projects/gap/cgi-bin/study.cgi?study_id=phs000372.v1.p1) and phenotypic collection is at the National Alzheimer’s Coordinating Center (https://www.alz.washington.edu/).

## ACKNOLEDEGMENTS

Dr. Lobach is supported by 5R21AG043710-02.

Genotyping is performed by Alzheimer’s Disease Genetics Consortium (ADGC), U01 AG032984, RC2 AG036528. Phenotypic collection is coordinated by the National Alzheimer’s Coordinating Center (NACC), U01 AG016976

Samples from the National Cell Repository for Alzheimer’s Disease (NCRAD), which receives government support under a cooperative agreement grant (U24 AG21886) awarded by the National Institute on Aging (NIA), were used in this study. We thank contributors who collected samples used in this study, as well as patients and their families, whose help and participation made this work possible;

Data for this study were prepared, archived, and distributed by the National Institute on Aging Alzheimer’s Disease Data Storage Site (NIAGADS) at the University of Pennsylvania (U24-AG041689-01)

We thank Ivan Belousov for help with the computations.

